# Rapid direct nucleic acid amplification test without RNA extraction for SARS-CoV-2 using a portable PCR thermocycler

**DOI:** 10.1101/2020.04.17.042366

**Authors:** Soon Keong Wee, Suppiah Paramalingam Sivalingam, Eric Peng Huat Yap

## Abstract

There is an ongoing worldwide coronavirus disease 2019 (Covid-19) pandemic caused by severe acute respiratory syndrome coronavirus 2 (SARS-CoV-2). At present, confirmatory diagnosis is by reverse transcription polymerase chain reaction (RT-PCR), typically taking several hours and requiring a molecular laboratory to perform. There is an urgent need for rapid, simplified and cost-effective detection methods. We have developed and analytically validated a protocol for direct rapid extraction-free PCR (DIRECT-PCR) detection of SARS-CoV-2 without the need for nucleic acid purification. As few as 6 RNA copies per reaction of viral nucleocapsid (N) gene from respiratory samples such as sputum and nasal exudate can be detected directly using our one-step inhibitor-resistant assay. The performance of this assay was validated on a commercially available portable PCR thermocycler. Viral lysis, reverse transcription, amplification and detection are achieved in a single-tube homogeneous reaction within 36 minutes. This minimized hands-on time, reduces turnaround-time for sample-to-result and obviates the need for RNA purification reagents. It could enable wider use of Covid-19 testing for diagnosis, screening and research in countries and regions where laboratory capabilities are limiting.

## Introduction

A novel coronavirus disease 2019 (Covid-19) caused by severe acute respiratory syndrome coronavirus 2 (SARS-CoV-2) was reported in Wuhan, China in December 2019 (1). The World Health Organization (WHO) has declared Covid-19 a pandemic, triggering various travel restrictions, border control, contact tracing, quarantine and social distancing measures in many countries (2). Despite these interventions, Covid-19 continues to spread, with 2 million cases reported globally (3), and even in countries with well-developed public health systems and aggressive implementation of measures (4). For instance, Singapore which reported its first imported case on 23^rd^ Jan 2020 and the first local transmission on 4 February 2020 (5, 6), has seen a ten-fold increase in cumulative cases in the past 25 days (7).

The first SARS-CoV-2 genome was published and deposited in NCBI database as Wuhan-Hu-1, GenBank accession number MN908947 on 14 January 2020 (8). This allowed several laboratories around the world, including our laboratory, to develop nucleic acid amplification tests (NAAT) to detect SARS-CoV-2 genetic materials (9). Currently, there are many quantitative RT-PCR (RT-qPCR) based NAAT being developed for SARS-CoV-2 (10). These target various viral genes including the nucleocapsid (N), polyprotein (ORF1ab), spike (S) and envelope (E) gene region of the positively stranded 29.9 kb RNA virus. These tests are widely used to screen suspected Covid-19 patients, returning travelers from outbreak areas, close contacts of cases, and healthy individuals who may be asymptomatic carriers of the virus (5, 11). Typically, respiratory samples such as sputum or nasal, throat, nasopharyngeal swabs collected consecutively over two days are tested to confirm a diagnosis or to confirm recovery. (5). Diagnostic testing is recognized as a rate-limiting step, and there is a global need to ramp up laboratory capacity, in both well-developed and low-income countries (4, 12). This is worsened by the possibility that asymptomatic individuals in the community could transmit infection, hence increasing the need to screen healthy individuals or those with mild symptoms (13–15).

Quantitative RT-PCR is being deployed as primary method for SARS-CoV-2 detection in research and hospital laboratories on account of its single-molecule sensitivity, ease of assay design and availability of reagents. However, there are several technical challenges. These tests require RNA extraction, followed by amplification and detection. Current state-of-art PCR typically require 70 minutes for RNA extraction and a further 90 minutes for amplification (15, 16). Highly trained technical staff, costly equipment (costing USD 20K - 50K) and facilities are required. These factors contribute to longer turnaround-time, costs of manpower, capital and consumables as well as risks of carryover contamination and biosafety risks when handling clinical samples. There is a limited supply of extraction reagents and test kits worldwide (16, 17). In addition, asymptomatic individuals have also led to pre-symptomatic transmission in the community which further increases demand for laboratory testing (13). These factors have motivated us to explore ways to simplify and shorten the protocols, without significant compromise to the high sensitivity and specificity of RT-qPCR. We therefore propose a novel method, which we termed direct rapid extraction-free PCR (DIRECT-PCR) for Covid-19 diagnosis.

PCR directly from crude samples without nucleic acid purification has been attempted before using inhibitor-resistant enzymes, modified buffers and additives in the mastermix. Our group (Sivalingam et al, unpublished) and others (18) have previously detected whole dengue virus in a single tube reaction containing serum and plasma in up to 8% (v/v). Direct amplification from samples has also been reported for the detection of other RNA viruses, including African Chikungunya virus (19), noroviruses (20), and bovine viral diarrhea virus (21) and from a variety of matrices, including serum, throat swab and faeces (22). However the presence of PCR inhibitors, such as mucin and proteins, poses a challenge for direct amplification from respiratory samples (23), and there are limited studies amplifying coronaviruses directly from such specimens. We have developed a DIRECT-PCR protocol using widely used and validated PCR primers, established its analytical performance with both DNA and RNA templates in respiratory samples, and transferred the protocols from benchtop to a portable thermocycler.

## Materials and Methods

### Samples and controls used for assay development

For biosafety purposes, synthetic nucleic acids rather than whole virus templates were used for the development of this assay. As SARS-CoV-2 is a positive stranded RNA coronavirus, we synthesized single stranded RNA (ssRNA) of the amplicon sequence (99 bp) for SARS-CoV-2 nucleocapsid (N) gene (Integrated DNA Technologies, San Diego). Plasmid DNA containing the N gene of SARS-CoV-2, MERS-CoV and SARS-CoV were also used as positive controls (2019-nCoV RUO Plasmid Controls, Integrated DNA Technologies, San Diego). ssRNA of human ribonuclease P (RP) gene amplicon sequence (65 bp) was synthesized for use as internal control (Integrated DNA Technologies, San Diego).

### Collection and processing of sputum and nasal exudate samples

Approximately 1 mL of SARS-CoV-2-negative sputum and nasal exudate from a healthy adult research team member was collected by hypersaline inhalation, and used for spike-in of ssRNA and plasmid controls. Briefly, the respiratory samples were mixed in 1:1 (v/v) ratio with Sputasol (Oxoid, England) and vortexed for 5 minutes to remove the viscosity to allow direct addition to the PCR reaction mix.

### Primer and probes

All primers and probes targeting SARS-CoV-2 ORF1ab and N genes were previously published by Chinese Center for Disease Control and Prevention, Beijing, China (24, 25). The human Ribonuclease P (RP) gene primer and probe were previously published by US Centers for Disease Control and Prevention (CDC) protocol and was added as an internal control to detect the presence of human RNA in the samples (26). All oligonucleotides, primers and probes were commercially purchased from Integrated DNA Technologies (San Diego).

### RT-qPCR and DIRECT-PCR

Quantitative RT-PCR was initially performed in monoplex single-tube reaction mixture using two different separate mastermixes. For standard RT-qPCR, PCR master mix containing Invitrogen SuperScript™ III Platinum™ One-Step RT-qPCR Kit (Life Technologies, USA) mastermix, 400 nM of forward and reverse primers, 200 nM of FAM-based probe, and 16 U of RNaseOUT™ Recombinant Ribonuclease Inhibitor (Life Technologies, USA) was used. For DIRECT-PCR, it was performed using a PCR enhancer and inhibitor-resistant enzymes (Direct One-Step S/P RT-qPCR TaqProbe Kit, VitaNavi Technology LLC, USA), supplemented with 400 nM of forward and reverse primers, 200 nM of FAM-based probe, and 16 U of RNaseOUT™ Recombinant Ribonuclease Inhibitor (Life Technologies, USA). 2 μL of ten-fold diluted RNA template in duplicates was added in a total volume of 20 μL. DNase/RNase-free water was used as the non-template control (NTC). All reactions were completed in a 96-well plate format (MicroAmp™ Fast Optical 96-Well Reaction Plate with Barcode, 0.1 mL). The RT-qPCR assays were performed under the following conditions: reverse transcription at 50°C for 15 minutes and initial denaturation at 95°C for 1 minute, 45 cycles of denaturation at 95°C for 10 seconds and annealing at 55°C for 45 seconds using a standard benchtop real-time thermocycler (StepOnePlus Real-Time PCR System, Applied Biosystems, USA). A specimen was considered positive if the amplification curve crossed the threshold line within 40 quantification cycle (C_q_ < 40).

### DIRECT-PCR of SARS-CoV-2 N gene in sputum and nasal exudate

All samples used for the spike-in experiments were freshly prepared. Briefly, 8.5 μL of sputum and nasal exudates sample mix with Sputasol were aliquoted. 0.5 μL (20 U) of RNaseOUT™ Recombinant Ribonuclease Inhibitor (Life Technologies, USA) was added to each aliquot and inverted to mix. Meantime, the RNA template was ten-fold serially diluted in DNase/RNase-free water to achieve 8 orders of magnitude. Next, 1 μL of diluted RNA template was spiked into the sputum and nasal exudate mix forming a total volume of 10 μL. Subsequently, 2 μL of RNA spiked-matrices containing 42.5% sputum and nasal exudate respectively were directly added into the 20 μL reaction volume of inhibitor-resistant PCR reaction mix containing a PCR enhancer and inhibitor-resistant enzymes (Direct One-Step S/P RT-qPCR TaqProbe Kit, VitaNavi Technology LLC, USA), forward and reverse primers, FAM-based probe, and RNaseOUT™ Recombinant Ribonuclease Inhibitor (Life Technologies, USA) as described above. RNA template in DNase/RNase-free water was used as positive control. A sputum only and nasal exudate only sample containing Sputasol were also added as a blank non-template control (NTC) for the respective matrices. The reaction mixture was incubated at 50°C for 15 minutes, denatured at 95°C for 1 minute, followed by 45 cycles of denaturation at 95°C for 10 seconds and annealing at 55°C for 45 seconds on a standard benchtop real-time thermocycler (StepOnePlus Real-Time PCR System, Applied Biosystems, USA).

The lowest limit of detection (LoD) was determined using SARS-CoV-2 N RNA template that was ten-fold serially diluted in SARS-CoV-2-negative sputum and nasal exudate samples. LoD was defined as the last dilution in which quantification cycle (C_q_) value could be detected in all replicates. The linear range correlation between the theoretical log copy number calculated from the concentration molarity of synthetic nucleic acids and C_q_ value was established applying a best-fit line to the data by linear regression analysis. PCR efficiency (E) was calculated from the slope of the linear equation.

### Optimization of Fast DIRECT-PCR assay

The DIRECT-PCR assay was modified in order to further reduce turnaround time and reagent cost. The final reaction volume was reduced from 20 μL to 10 μL, while reducing the number of cycles from 45 to 40, reducing reverse transcription (RT) step from 15 minutes to 5 minutes, reducing initial denaturation from 1 minute to 30 seconds and reducing annealing duration from 45 seconds to 15 seconds. Ten-fold serially diluted RNA spike-in matrices described above were used in duplicates to evaluate the modified assay to determine a fast protocol for detection of SARS-CoV-2. Briefly, an aliquot of 1 μL of template was added in the 10 μL PCR reaction mix mentioned above. Similarly, the plasmid controls of N gene from SARS-CoV-2, MERS-CoV and SARS-CoV were ten-fold serially diluted in DNase/RNase-free water to achieve 6 orders of magnitude respectively. Subsequently, 1 μL of diluted plasmid template was spiked into the sputum and nasal exudate mix containing Sputasol and RNaseOUT forming a total volume of 10 μL as described above. The internal control using human Ribonuclease P (RP) gene was evaluated by the DIRECT-PCR of human RP in the sputum and nasal exudate using the assay described. Serially diluted human RP RNA template in water was used as the positive control. MERS-Cov and SARS-Cov plasmid controls were added as negative controls. The amplification was performed on a standard benchtop real-time thermocycler (StepOnePlus Real-Time PCR System, Applied Biosystems, USA).

The detailed protocol of DIRECT-PCR for SARS-CoV-2 is available as Supplementary Information (Supplementary S1).

### Performance of portable real-time thermocycler

DIRECT-PCR assays were validated on the portable thermocycler (MyGo Mini, IT-IS Life Science Ltd, Ireland) using the same monoplex single-tube protocols described above. The portable thermocycler is lightweight at less than 2 kg with a dimension of 12 cm (width) by 12 cm (depth) by 16 cm (height). It can perform up to 16 reactions using standard 0.1 mL clear qPCR tubes with 10 to 100 μL reaction volume each. The performance of this portable qPCR thermocycler was evaluated by comparing the LoD and amplification efficiency (E) of DIRECT-PCR assays.

### Statistical Analysis

The limit of detection (LoD) was determined by plotting quantification cycle (C_q_) against log_10_ copy number concentration. Correlation coefficient (R^2^) was calculated by linear regression analysis. Amplification efficiency (E) was calculated from the slope of the log-linear curve using the given equation: E = −1+10^(−1/slope)^. The slopes and intercepts of the linear regression lines were tested for statistical significance using Analysis of Covariance (ANCOVA) (GraphPad Prism 6). Repeatability and reproducibility of the RT-qPCR assay was determined by analysing the mean values and standard deviations of C_q_ values.

## Results

### Determination of LoD and amplification efficiency (E)

Using synthetic RNA template of the SARS-CoV-2 N gene target, the LoD of standard RT-qPCR was assessed to be 120 RNA copies per reaction with a C_q_ mean value of 40.67 ± 0.29. However, no amplification was observed when RNA was spiked in sputum and nasal exudate using this standard mastermix. Using a PCR inhibitor-resistant mastermix, the LoD of DIRECT-PCR was determined to be 120 RNA copies per reaction with a C_q_ mean value of 38.48 ± 0.57 at the 7^th^ order of magnitude (Table 1). The lowest C_q_ was observed at 17.37 ± 0.04 which corresponded to 1.2 × 10^8^ RNA copies per reaction (Fig. 1A). The DIRECT-PCR assay was completed in 72 minutes on the benchtop thermocycler. The control assay has amplification efficiency (E) of 84.94% with a correlation coefficient (R^2^) of 0.9884 (Fig. 1D). No amplification was observed for the NTC.

**Table 1.** Limit of detection (LoD) in copies per PCR reaction volume and PCR efficiency of DIRECT-PCR for SARS-CoV-2 on benchtop and portable thermocyclers.

| PCR | Mastermix volume per reaction (μL) | Mastermix used | Template | Matrix | Benchtop Thermocycler |  |  | Portable Thermocycler |  |  |
| --- | --- | --- | --- | --- | --- | --- | --- | --- | --- | --- |
|  |  |  |  |  | LOD (C<sub>q</sub> Mean ± S.D) | PCR Efficiency (%) | R<sup>2</sup> | LOD (C<sub>q</sub> Mean ± S.D) | PCR Efficiency (%) | R<sup>2</sup> |
| RT-qPCR | 20 | Invitrogen SuperScript™ III Platinum™ One-Step RT-qPCR Kit | ssRNA | Water<br>Sputum<br>Nasal Exudate | 120<br>(40.67 ± 0.29)<br>No amplification<br>No amplification | 84.30 | 0.9994 | 120<br>(38.70 ± 0.10)<br>No amplification<br>No amplification | 86.26 | 0.9985 |
| DIRECT-PCR | 20 | VitaNavi Direct One-Step S/P RT-qPCR TaqProbe Kit | ssRNA | Water<br>Sputum<br>Nasal Exudate | 120<br>(38.48 ± 0.57)<br>12<br>(38.79^)<br>12<br>(38.72^) | 84.94<br>82.62<br>81.27 | 0.9884<br>0.9944<br>0.9986 | 120<br>(36.99^)<br>12<br>(38.10^)<br>1200<br>(36.47 ± 0.23) | 88.35<br>88.64<br>77.45 | 0.9729<br>0.9924<br>0.9976 |
| Fast DIRECT-PCR | 10 | VitaNavi Direct One-Step S/P RT-qPCR TaqProbe Kit | ssRNA | Water<br>Sputum<br>Nasal Exudate | 600<br>(39.42^)<br>6<br>(39.28^)<br>60<br>(39.34^) | 81.72<br>76.03<br>69.23 | 0.9859<br>0.9824<br>0.9865 | 600<br>(36.63^)<br>600<br>(36.25 ± 0.46)<br>60<br>(36.90^) | 89.37<br>85.52<br>81.08 | 0.9639<br>0.9784<br>0.9775 |
|  |  |  | Plasmid | Water<br>Sputum<br>Nasal Exudate | 2<br>(39.75^)<br>2<br>(38.93^)<br>20<br>(38.02 ± 0.55) | 119.25<br>101.91<br>96.72 | 0.9896<br>0.9932<br>0.9784 | 20<br>(36.56 ± 0.26)<br>20<br>(35.40 ± 1.35)<br>20<br>(36.66 ± 0.08) | 113.17<br>100.25<br>107.33 | 0.9669<br>0.9756<br>0.9931 |

^ one out of two duplicates positive

**Figure 1.**
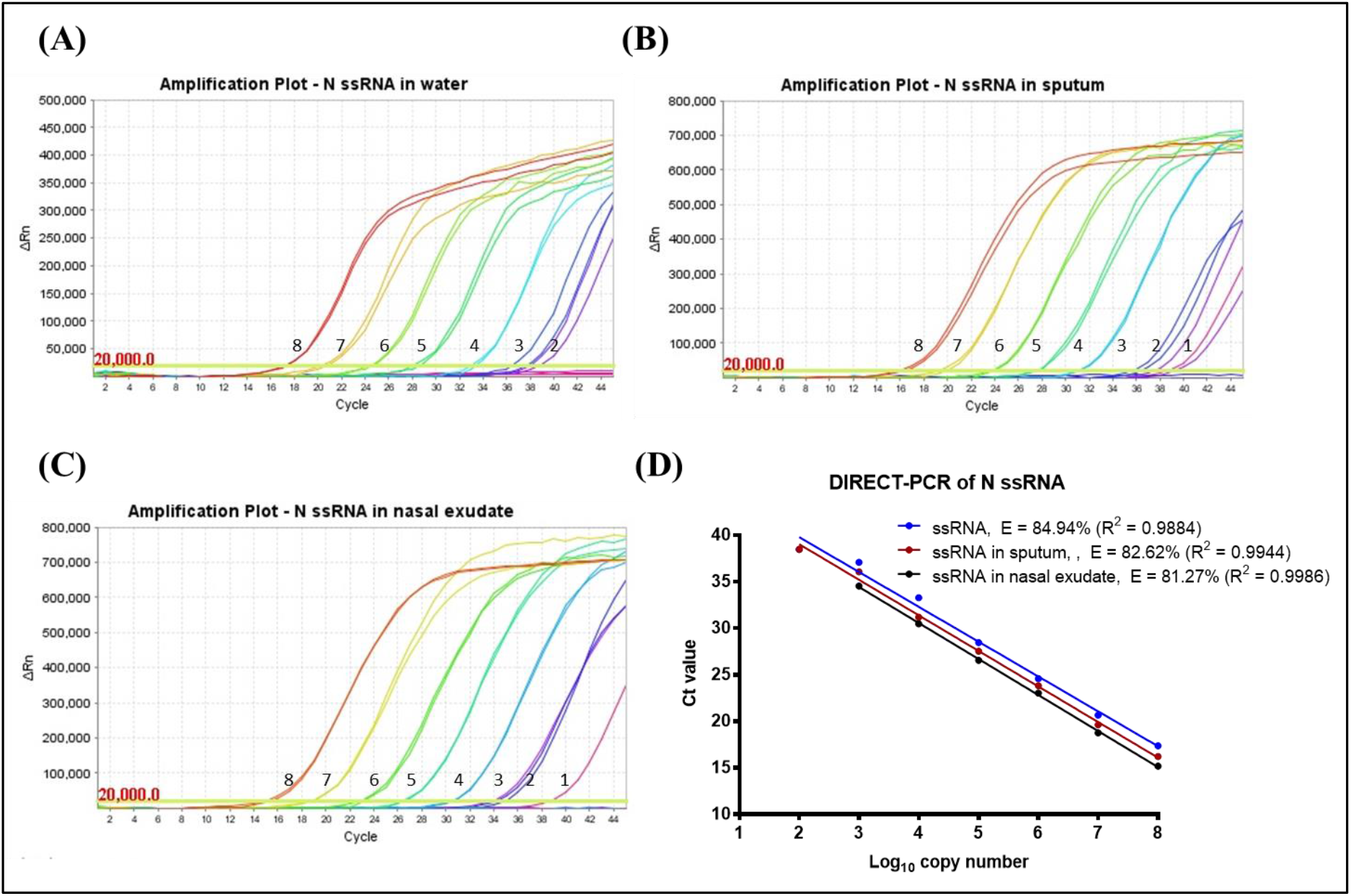
Amplification plot of SARS-CoV-2 N gene ssRNA control in (A) water, (B) spiked in sputum, (C) spiked in nasal exudate conducted in the same thermocycler run. Numbers indicated the log_10_ copy number of the template present. No amplification was observed in the NTC. (D) Comparison of DIRECT-PCR assay amplification of SARS-CoV-2 N gene ssRNA control (blue line), N gene spiked in sputum (red line) and N gene spiked in nasal exudate (black line). Templates were ten-fold diluted in 8 orders of magnitude. 2 μL of template was used in 20 μL of PCR mastermix on the benchtop thermocycler. The amplification efficiency (E) was determined by plotting of mean C_q_ values against log_10_ copy number calculated using theoretical molarity of templates.

Next, the DIRECT-PCR assay was evaluated using RNA spiked in sputum and nasal exudate matrices. The amplification of spiked RNA with PCR inhibition tolerance was observed with the concentration of sputum and nasal exudate at 4.25% (v/v) in the PCR reaction. In RNA spiked sputum samples, the sensitivity was greater than the control at 12 RNA copies per reaction with a C_q_ of 38.79 (Table 1). The lowest C_q_ was observed at 16.22 ± 0.24 which corresponded to 1.2 × 10^8^ RNA copies per reaction (Fig. 1B). When RNA was spiked in nasal exudate, we found the LoD of 12 RNA copies per reaction with a C_q_ of 38.72 (Table 1). These LoDs were achieved in one out of two duplicates tested. The lowest C_q_ in nasal exudate was observed at 15.18 ± 0.36 which corresponded to 1.2 × 10^8^ RNA copies per reaction (Fig. 1C). The PCR efficiency revealed lower amplification efficiency (E) when spiked into sputum (82.62%) and nasal exudate (81.27%) compared to control while the correlation coefficient (R^2^) at 0.9944 and 0.9986 for sputum and nasal exudate respectively. The linear regression lines had similar slopes (p-value = 0.6503) while the intercepts were significantly different (p-value < 0.0001) (Fig. 1D).

### Evaluation of Fast DIRECT-PCR assay

To reduce the duration and cost of the DIRECT-PCR assay, we modified the total reaction volume to 10 μL and duration of cycling conditions with the addition of 1 μL template. Amplification of RNA template using the fast DIRECT-PCR demonstrated detection at 600 RNA copies per reaction (C_q_ of 39.42) in contrast to 120 RNA copies per reaction (C_q_ of 38.48 ± 0.57) detected on the DIRECT-PCR assay. This fast DIRECT-PCR was completed in 36 minutes as compared to 72 minutes for the DIRECT-PCR assay. The LoD was similar at 6 (C_q_ of 39.28) and 60 RNA copies per reaction (C_q_ of 39.34) when spiked into sputum and nasal exudate samples respectively (Table 1). Using positive control plasmids encoding the N gene of SARS-CoV-2, the LoD was found to be 2 copies per reaction (C_q_ of 39.75) using the benchtop thermocycler. LoD of 2 and 20 copies per reaction was observed when the plasmids were spiked into sputum and nasal exudate samples with C_q_ of 38.93 and C_q_ mean of 38.02 ± 0.55 respectively over four orders of magnitude (Fig. 2). No amplification was observed for the negative controls using MERS-CoV and SARS-CoV plasmids. Human RP primers were used as an internal control to amplify and detect the presence of human RNA present in the crude samples (data not shown). Using crude sputum and nasal exudate directly, the mean C_q_ values for human RP was 26.12 ± 0.24 and 27.76 ± 0.81 respectively on the benchtop thermocycler.

**Figure 2.**
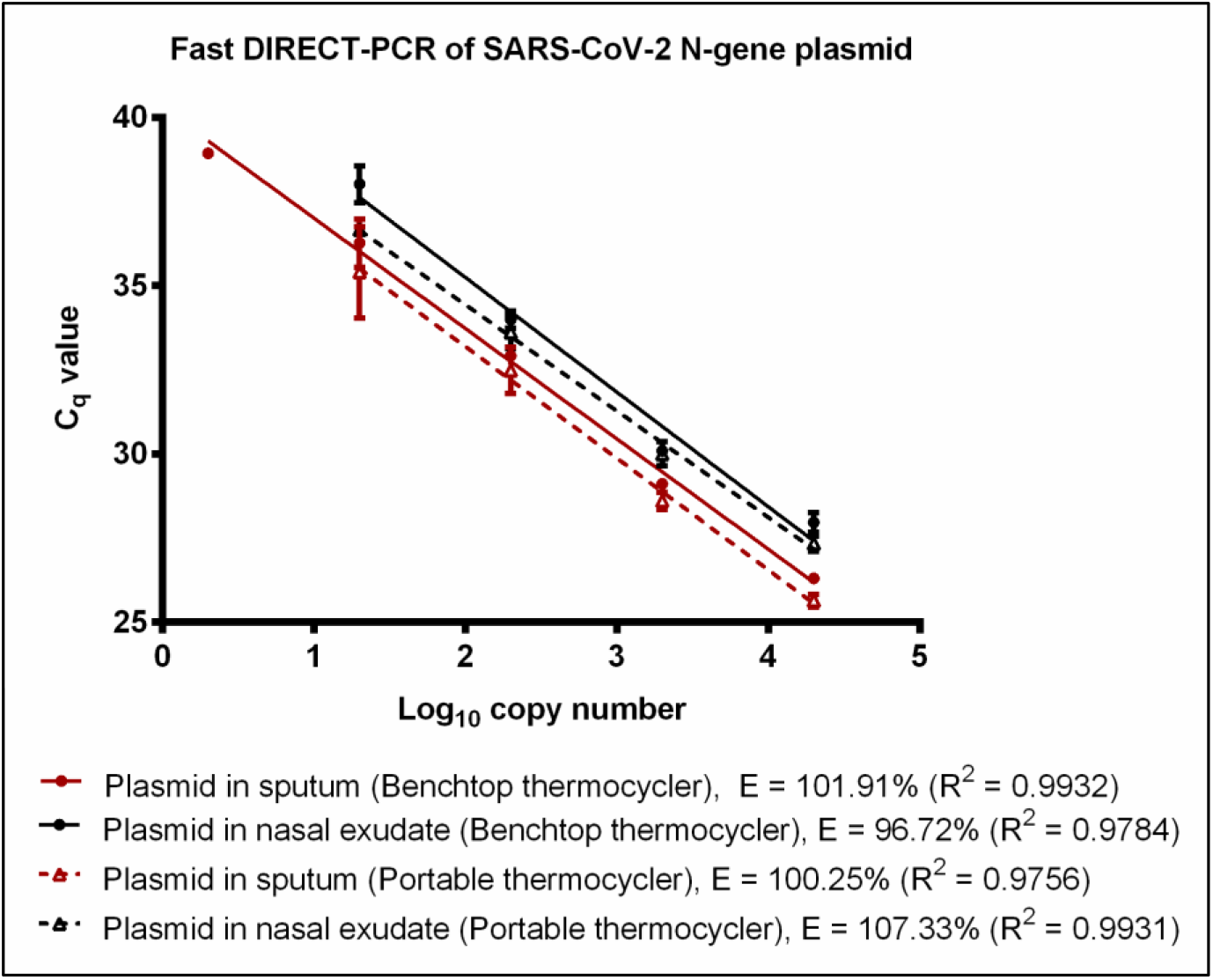
Fast DIRECT-PCR assay amplification efficiency of SARS-CoV-2 N gene plasmid spiked in sputum (red line) and N gene plasmid spiked in nasal exudate (black line) using the benchtop thermocycler (solid line) and the portable thermocycler (dotted line). Templates were ten-fold diluted in 6 orders of magnitude. 1 μL of template was used in 10 μL of PCR mastermix.

### Performance of portable thermocycler

A commercially available portable qPCR thermocycler was validated by comparing to the benchtop thermocycler using the fast DIRECT-PCR assay of RNA and plasmid templates spiked in sputum and nasal exudate. The amplification efficiency of RNA control showed that there was no significant difference (p-value = 0.4344) between benchtop and portable thermocycler. Both thermocyclers had similar LoD at 600 RNA copies per reaction. When using RNA spike-in sputum, the portable thermocycler had LoD of 600 RNA copies per reaction (C_q_ of 36.25 ± 0.46) compared to 6 RNA copies per reaction when using the benchtop thermocycler (Table 1). On the other hand, both thermocyclers had the same LoD of 60 RNA copies per reaction when using RNA spike-in nasal exudate (Table 1). Using plasmid controls, there was no significant difference (p-value = 0.7799) in amplification efficiency between both thermocyclers (Fig. 2). Next, using plasmids encoding the N gene of SARS-CoV-2 as control, the LoD was found to be 2 and 20 copies per reaction when using plasmid template and plasmid spike-in sputum for benchtop and portable thermocycler respectively. Both thermocyclers had similar sensitivity of 20 copies per reaction when using plasmid spike-in nasal exudate (Table 1). In terms of duration, the reaction took 49 minutes on the portable thermocycler and 36 minutes on benchtop thermocycler.

## Discussion

Quantitative RT-PCR of SARS-CoV-2 RNA is currently the method of choice for the diagnosis of Covid-19. In our DIRECT-PCR method, we used inhibitor-resistant enzymes and reagents to eliminate the RNA extraction step. The use of synthetic RNA as template obviated the need to handle viral nucleic acids. With fast DIRECT-PCR protocol, we achieved further reduction in assay time by optimizing the thermocycling conditions. We also validated the analytical performance of DIRECT-PCR on spiked crude samples and demonstrated its use on a portable platform.

Current RT-qPCR for SARS-CoV-2 involves RNA purification as part of the pre-PCR sample preparation procedure and at least four to six hours are needed for time-to-results in most laboratories (16). When the cycling conditions were optimized in our fast DIRECT-PCR with small reaction volumes (10 μL), and shorter RT and annealing durations, we were able to amplify SARS-CoV-2 in less than an hour (Fig. 3). Furthermore, there are cost savings on nucleic acid extraction kits, and the availability of reagents themselves may be limiting when laboratory capacity is ramped up and demand is increased globally. Moreover, DIRECT-PCR is a single-tube homogeneous reaction that reduces hands-on time and biosafety risk for laboratory personnel, as well as the likelihood for carryover contamination. One caveat however for fast DIRECT-PCR of samples with low viral load is that of sampling error, since only 1 μL of sample is used. Nucleic acid extractions, on the other hand, serve to concentrate RNA from typical sample volumes of 150-300 μL, although their yield can also be low and variable. Hence, where samples are expected to have low counts near to the limit of detection, DIRECT-PCR with larger volume reactions (25 μL) to include higher template volume may be necessary to reduce risk of false negatives.

**Figure 3.**
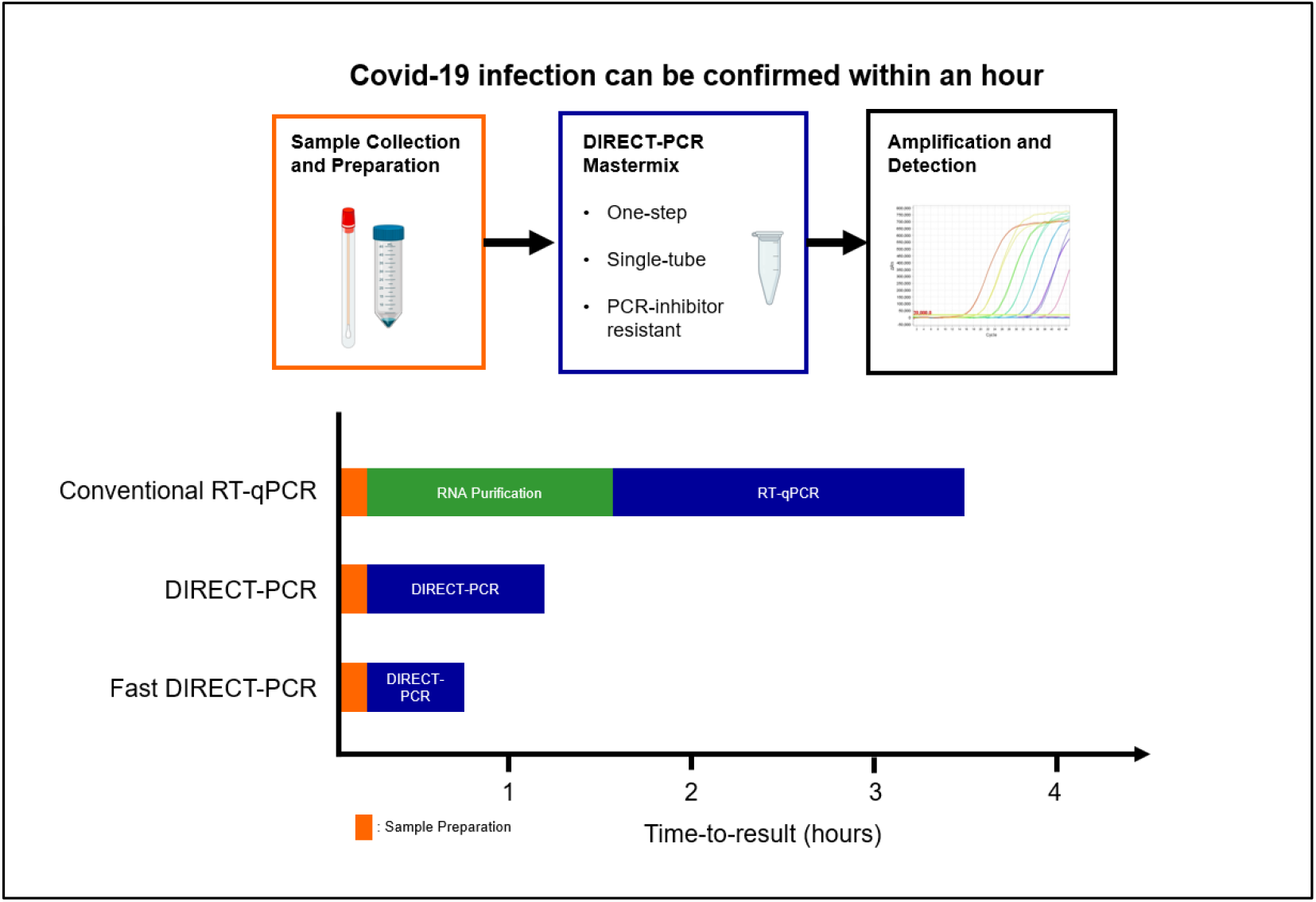
Workflow of fast DIRECT-PCR protocol for the direct detection of SARS-CoV-2. Covid-19 infection can be confirmed in less than an hour after sample collection and addition to mastermix for amplification and detection.

Reduction in amplification efficiency is a common concern in DIRECT-PCR from crude samples (e.g. respiratory samples, blood and serum). The presence of PCR inhibitors can decrease the sensitivity and accuracy of pathogen detection through interfering with polymerase activity, degradation of nucleic acids and efficient cell lysis (23). A variety of methods have been developed to overcome such inhibition, including inhibitor tolerant polymerases, additives and buffer modification. In our study, we used one commercially available formulation that tolerated PCR inhibition in sputum and nasal exudates as well as blood/serum/urine (data not shown), and it is not unlikely that other formulations could be used as well in our study. While amplification directly from sputum and nasal exudate reduced the efficiency of PCR compared to water controls, there was no net effect on the threshold of detection.

Nasopharyngeal swabs and sputum have been suggested to be effective clinical samples for diagnosis (27, 28), and we have assessed direct PCR using these matrices. It should be feasible to extend this to other common samples used for Covid-19 testing, such as swabs of the throat and endotracheal tubes, and bronchoalveolar lavage fluid. While we have focused our DIRECT-PCR protocols on commonly used primer-probe sets developed by the China CDC and US CDC that target the SARS-CoV-2 N gene as well as ORF1ab gene and the internet control RP (data not shown), we expect direct PCR to be similarly useful for other viral primer targets and PCR amplicons developed for Covid-19 (8, 9), as well as for multiplexed amplification of several targets in the same tube. Nevertheless, there may be other inhibitors of PCR that were not present in our samples that we tested, and clinical validation with a larger series is needed to exclude this possibility.

Quantitation of viral nucleic acid in samples by RT-qPCR has been used in studies to correlate viral load with severity, prognosis and transmissibility. The C_q_ values in patients’ samples can range from 19 to late cycles close to 40 during the course of infection. The nasal viral load has been shown to peak within days of symptom onset according to small study of throat and nasal swabs from 17 patients (29). Using RNA as template, our DIRECT-PCR assay has a dynamic range of over 7 orders of magnitude with mean C_q_ range from 17 to 38, indicating it is reproducibly quantitative over a wide range, including very high viral titers. This method could be useful in future studies, such as to assess viral survivability, decontamination methods and preventive intervention.

We have observed, even in spiked water samples, that amplification efficiencies and the C_q_ values obtained at the limit of detection are dependent on factors such as the reagent mastermix, the model of thermocycler, the reaction volumes and the cycling protocols used (Table 1). This variability would be even more significant in a clinical setting, with the added confounding factors such as type of sample, sampling method, dilution in transport buffers, duration and conditions of storage, and the reagents and protocols for RNA extraction, that are difficult to standardize across healthcare settings and laboratories. While removing the RNA extraction step could help to reduce this variability in RNA yield and quality, variability in sample type and sampling procedure still renders the entire assay non-quantitative. Hence, we caution over-interpretation of C_q_ values, especially in the absence of in-house quality control and calibration data for each combination of protocol, reagents and thermocycler. These technical factors may also partly explain some of the false negatives and temporal variability in viral RNA shedding reported in various studies, that use fixed criteria C_q_ < 37 for positive, C_q_ between 37 to 40 as inconclusive for repeat testing and C_q_ > 40 as negative. In fact, our data demonstrates that as viral RNA concentrations approach the threshold of detection, our assay LoD has C_q_ mean value of > 37 and may exceed 40 (Table 1). Since negative controls in these probe-based assays remain consistently negative, it should be possible to report C_q_ > 37 as positive. Hence to achieve better standardization of quantitative results, we propose that instead of reporting C_q_, laboratories may report the equivalent viral copy number in per unit sample volume, derived from running serial dilutions of viral or RNA controls in-house.

Our DIRECT-PCR assay has been demonstrated on a commercially available portable thermocycler that weighs less than 2 kg, potentially allowing PCR to be performed outside molecular biology laboratories, in mobile laboratories, and in low resource settings. There are many real-time fieldable thermocyclers available, that are smaller, portable and less costly than benchtop ones, and three of these we have evaluated perform similar to benchtop ones (data not shown). While the optical detection modules and their fluorescence sensitivity vary between designs, the overall sensitivity of our assay was similar in both types of thermocyclers tested (Table 1). Thermocycling parameters could also be optimized for rapidity (Fig. 3). As the need to transport samples to central laboratories could prolong the availability of test results to beyond 24 hours (16), use of portable thermocyclers, coupled with appropriate training and quality control procedures, could allow the use of RT-qPCR nearer to the patients and in primary healthcare settings. Such rapid point-of-care (POC) tests for use at community level were identified by a WHO expert group as a key research priority (30). As the need to transport samples to central laboratories could prolong the availability of test results to beyond 24 hours (16), use of portable thermocyclers, coupled with appropriate training and quality control procedures, could allow the use of RT-qPCR nearer to the patients and in primary healthcare settings (12).

Though it is unlikely that DIRECT-PCR would completely replace conventional RNA extraction with RT-qPCR in large laboratories as a confirmatory diagnosis, the trade-off in reliability for portability, ease of use and cost-effectiveness could be useful for patient screening and public health surveillance and in settings with limited laboratory resources and facilities (4). DIRECT-PCR for Covid-19 could address the specific requirements and challenges of delivering molecular genetic testing at the POC setting. POC diagnostics could play an important role in the detection, diagnosis and control of this pandemic (17), such as for more timely diagnosis in primary healthcare facilities, and to complement real-time fever surveillance and screening. Several technologies have been proposed and used to enable molecular testing at POC, including isothermal amplification methods that do not require such thermocyclers, like loop-mediated isothermal amplification (LAMP) (12). However, given the wide-spread validation of existing SARS-CoV-2 PCR primer/probe sets, as well as the likely need to modify amplification primers and probes as the viral sequences evolve in the future, we argue that performing PCR using our DIRECT-PCR method is a more generic and adaptable approach.

## Conclusion

Direct amplification of SARS-CoV-2 viral RNA from samples without RNA purification has been developed, reducing hands-on-time, time-to-results, and costs. Viral lysis, reverse transcription, amplification and detection are achieved in a single-tube homogeneous reaction taking less than an hour, and they can be performed on a portable thermocycler. Analytical validation was performed with sputum and nasal exudate. The DIRECT-PCR assay has a high sensitivity of 6 RNA copies per reaction and is quantitative over a dynamic range of 7 orders of magnitude. This method may be useful during the current global Covid-19 pandemic in situations where resources are constrained or where timely results are needed.

## Acknowledgment

This research is supported by a Start-Up Grant from Lee Kong Chian School of Medicine, Nanyang Technological University Singapore.

## Contributions

S. K. W. and S. P. S. contributed equally to this work. S. K. W., S. P. S. and E. P. H. Y. confirmed they have contributed to the intellectual content of this paper and have met the following 3 requirements: (a) significant contributions to the conception and design, acquisition of data, or analysis and interpretation of data; (b) drafting or revising the article for intellectual content; and (c) final approval of the published article.

