## Supplementary Information S1 for "Rapid direct nucleic acid amplification test without RNA extraction for SARS-CoV-2 using a portable PCR thermocycler"

**Protocol for DIRECT-PCR of SARS-CoV-2 from crude respiratory samples**

**Background**
This protocol describes the single-tube one-step DIRECT-PCR protocol of SARS-CoV-2 from crude respiratory samples. It details the preparation of DIRECT-PCR mastermix and sample processing.

**Materials and Methods**

Table S1. Consumables used for the direct detection of SARS-CoV-2.

| Reagents and Consumables | Storage Condition |
|---|---|
| 10 µM Primers and Probes | -30°C freezer |
| Invitrogen™ RNaseOUT™ Recombinant Ribonuclease Inhibitor (5000 U, Specific activity: 40 U/µL,<br>ThermoFisher Catalog 10777-019) | -30°C freezer |
| Molecular grade DNase/RNase-free water | 4°C fridge |
| VitaNavi Technology Direct One-Step S/P RT-qPCR TaqProbe Kits (No Reference Dye)<br>• VitaNavi 2X S/P RT-qPCR TaqProbe Mix<br>• VitaNavi S/P RT-qPCR Pols | -30°C fridge |
| General Consumables:<br>• Micropipette<br>• Aerosol-resistant tips<br>• Centrifuge<br>• Vortex<br>• Sputasol (Oxoid)<br>• 1.5 ml Microcentrifuge Tube<br>• qPCR plate<br>• qPCR plate optical adhesive film<br>• PCR 8-strip tubes and caps | - |

**Primers and Probes**

Table S2. Primers and Probes used for the direct detection of SARS-CoV-2.

| Target | Primer Name | Sequence (5' to 3') | Amplicon size | Reference |
|---|---|---|---|---|
| Human Ribonuclease P control | RP-F | AGATTTGGACCTGCGAGCG | 65 bp | (CDC, 2020) |
| | RP-R | GAGCGGCTGTCTCCACAAGT | | |
| | RP-Probe | FAM–TTCTGACCTGAAGGCTCTGCGCG–BHQ1 | | |
| SARS-CoV-2 ORF1ab | SARS-CoV-2_ORF1ab-F | CCCTGTGGGTTTTACACTTAA | 119 bp | (C. CDC, 2020; Wang et al., 2020) |
| | SARS-CoV-2_ORF1ab-R | ACGATTGTGCATCAGCTGA | | |
| | SARS-CoV-2_ORF1ab-probe | FAM-CCGTCTGCGGTATGTGGAAAGGTTATGG-BHQ1 | | |
| SARS-CoV-2 N | SARS-CoV-2_N-F | GGGGAACTTCTCCTGCTAGAAT | 99 bp | |
| | SARS-CoV-2_N-R | CAGACATTTTGCTCTCAAGCTG | | |
| | SARS-CoV-2_N-Probe | FAM-TTGCTGCTGCTTGACAGATT-TAMRA | | |

**Preparation of Mastermix Area**

1. Setup separate physical areas for mastermix reagent preparation, RNA sample preparation and post-amplification area.
2. Disinfect all work surfaces with appropriate disinfectant.
3. Establish a unidirectional workflow to reduce contamination.
4. Do not mix materials or equipment from the sample preparation area and reagent preparation area.
5. Use separate set of pipettes are available for use in each area.
6. Use aerosol resistant tips to prevent aerosols and reduce contamination.
7. Always maintain ice box or cooler box for use to prepare reagents and samples in cold chain.

**PCR Mastermix Preparation**

Table S3. Concentration and volume of PCR mastermix reagents for 10 µl reaction

| Reagent | Final Conc. | Volume (µl) per rxn |
|---|---|---|
| VitaNavi 2X S/P RT-qPCR TaqProbe Mix | 1X | 5.0 |
| VitaNavi S/P RT-qPCR Pols | 1X | 0.5 |
| Invitrogen™ RNaseOUT™ Recombinant Ribonuclease Inhibitor | 16 U | 0.2 |
| 10 µM Forward primer | 400 nM | 0.4 |
| 10 µM Reverse primer | 400 nM | 0.4 |
| 10 µM FAM-Probe | 200 nM | 0.2 |
| Template | - | 1.0 to 2.0 |
| Molecular grade $H_20$ | - | Adjust the volume accordingly |
| Total | - | 10.0 |

1. Bring all reagents on ice (Table S3).
2. Flick the reagents to mix before use.
3. Include PCR positive template control sample, and a non-template control (NTC) in each run
4. Prepare separate MasterMix (MM) for each primer set (SARS-CoV-2 N gene, SARS-CoV-2 ORF1ab gene and human RP gene)
5. Label 1.5 mL microcentrifuge tube as MasterMix.
6. Add all the components in the following order:
    a. Add water first
    b. Add forward and reverse primers
    c. Add TaqProbe mix buffer
    d. Add probes
    e. Add RNaseOUT
    f. Add VitaNavi S/P RT-qPCR Pols last.
7. Mix the MasterMix by inverting the tube at least 10 times.
8. Aliquot MasterMix equally into qPCR plate/tubes (e.g. for 1 µL template, add 9 µL of MasterMix)
9. Cover the plate with aluminium foil to minimise probe exposure to light.

**Clinical Sample Preparation**

1. All clinical specimens must be carried out in a Class II Biological Safety Cabinet (BSC) in Biosafety Level-2 (BSL-2+) Plus Laboratory in accordance to local authority guidelines.
1. Setup Class II Biological Safety Cabinet (BSC).
2. Heat inactivate respiratory samples such as sputum or nasal, throat, nasopharyngeal swabs at 60°C for 10 minutes immediately upon receiving them.
3. Aliquot 1 mL of the samples into a sterile tube.
4. Pre-prepare working concentration Sputusol by adding entire vial content (7.5 mL) to 92.5 mL of sterile water accordingly to manufacturer's instruction.
5. Add 1 mL of Sputasol (in working concentration) to mix with the sample. The ratio of sample : sputasol is 1:1 (v/v).
6. Vortex for 5 minutes.

**Addition of templates into MasterMix**

1. Add 1 to 2 µL of samples into the MasterMix well without introducing bubbles.
   *Note: This brings the final PCR reaction volume to 10 µL.*
2. Add 1 µL of RNA positive control sample into the positive control well.
3. Add 1 µL of non-template control (NTC) sample into the negative control well.
4. Seal the qPCR strip-tube with the optical cap / qPCR plate with the optical adhesive film.
5. Briefly centrifuge (spin-down) the qPCR mix and ensure that there are no bubbles.

**PCR Protocol**

1. Setup the qPCR Machine according to manufacturer's instructions.
   a. Example: Comparative CT mode, 96-well, Taqman, fast ramp rate, FAM-Probe, No quencher
2. Setup the dRT-qPCR run condition as follows:
   a. Reverse Transcription: 50°C – 5 minutes
   b. Initial denaturation: 95°C - 30s
   c. Cycling stage: 40 x (95°C - 10s + 55°C - 15s)

**Analysis of Results**

1. After completion of the run, save and analyze the experimental data following the instrument manufacturer's instructions.
2. Analyses should be performed separately for each target using a manual threshold setting.
3. Thresholds should be adjusted to fall within exponential phase of the fluorescence curves and above any background signal. The procedure chosen for setting the threshold should be used consistently.
4. Negative for NTCs (with water) must be blank.
5. Positive controls must be positive with < 40 $C_q$ value.
6. Internal RNase P controls for clinical samples must be positive with < 40 $C_q$ value.
7. If SARS-CoV-2 N gene and ORF1ab gene crosses the cycle threshold line within 40 $C_q$, the samples will be considered positive for Covid-19 infection.
